# Precision mapping of the mouse brain metabolome

**DOI:** 10.1101/2020.12.28.424544

**Authors:** Huanhuan Pang, Jun-Liszt Li, Xiao-Ling Hu, Fei Chen, Xiaofei Gao, Lauren G. Zacharias, Feng Cai, Ralph J. DeBerardinis, Wenzhi Sun, Zeping Hu, Woo-ping Ge

## Abstract

Metabolism is physiologically fundamental to a biological system. Understanding brain metabolism is critical for our comprehensive knowledge of brain function in health and disease. Combining a microarray collection system with targeted metabolomics analysis, here we performed precision mapping of the metabolome in the mouse brain and created maps for 79 metabolites with a resolution of 0.125mm^3^ per pixel (i.e., brain subregion). The metabolome atlas provides researchers with a useful resource to interpret the vulnerability of specific brain regions to various disease-relevant metabolic perturbations.

---

Many diseases, including those manifesting in the brain, heart, muscle, liver and other organs, involve localized metabolic perturbations that interfere with physiological tissue function and contribute to pathogenesis(*1–3*). The severity of many neurological diseases varies with the location of the associated lesion(s) in the brain. For example, the hippocampus and entorhinal cortex are the first brain regions to be affected by amyloid and neurofibrillary pathology in patients with Alzheimer’s disease(*4*). The heterogeneity of metabolism in different brain regions may be due to the expression pattern of metabolic enzymes or solute carriers (SLC) varying in the neurons or glial cells(*5, 6*). These distinct metabolic differences can cause variations in neuronal survival or degeneration under pathological conditions in the brain. Precision mapping of the brain metabolome would help elucidate the physio-pathologies and underlying mechanisms of these neurological diseases.

Although regional metabolomic analyses have been used to characterize the brain(*7, 8*), a precise and high-resolution in situ metabolic database of the whole brain is still lacking. Mass spectrometry imaging (MSI), including Matrix-assisted laser desorption/ionization (MALDI), SIMS and DESI(*9*) allows the direct analysis of measurement of both abundance and distribution of protein, lipids(*10*), peptides and even metabolites from different mammalian tissues or surgical samples. In particular, MALDI has enabled *in situ* mapping of some neurotransmitters in the brain(*11–13*), and has made impressive advances in the field(*14, 15*). However, measurements using MSI can vary substantially due to factors such as thin slice preparation, deposition manipulation, matrix selection, as well as desorption and ionization(*16*). Managing all of these variables requires exceptional skill, as a result largely limiting MALDI’s application for precise and large-scale mapping of the brain metabolome(*17*). In addition, MSI of many metabolites with molecular weight <1,000 DA is much more challenging than MSI of proteins, lipids and peptides due to multiple factors: matrix compounds introduce interference in the mass range of small molecules; metabolites have rapid turnover rates; and some metabolites have low *in vivo* concentrations(*18*). In this study, we developed a strategy to divide a brain section into ~300 subregions. Using targeted metabolomic analysis, we generated a two-dimensional (2-D) metabolic atlas of a hemisphere of a mouse brain for a large number of metabolites.

To overcome the challenge of achieving even tissue collection from a small brain region, we developed a microarray collection system to dissect out tiny regions of pre-defined size from freshly-isolated brain slices (**Fig. 1A–C, fig. S1**). The collection system was made of stainless steel tubes with different outer diameters (OD, ranging from 0.008 in to 0.0390 in, i.e., 203 μm to 991 μm). The area of one typical mouse brain section including the hippocampus is about 80 mm^2^, so the OD of a micro-collection tube determines the resolution of brain metabolome mapping (resolution = the number of pixels in one brain slice = 80 mm^2^/OD^2^). For 203 μm tubes, one 80 mm^2^ brain section can be divided into 1,941 fractions (i.e., pixels). The inter diameter (ID) of a tube determines the number of cells that we collect per sample. For a tube with OD 203 μm and ID 165 μm used for tissue collection from a brain slice (500 μm thickness, ~40 mg), we obtained ~10.7 μg brain tissue per sample. Given that there are ~109 million cells in one mouse brain (~0.4 g/brain)(*19*), the tissue per sample equated to roughly 2,916 cells that we collected from one of these microtubes. Although this method allowed us to detect 30–40 metabolites per sample, we modified the approach to increase the metabolite yield by creating micro-collection tubes with a larger diameter and thinner tube wall for our subsequent mapping experiments (OD, 0.0200 in; ID, 0.0165 in, i.e., OD, 508 μm; ID 419 μm). With this type of tube for collection, we obtained 68.9 μg brain tissue per tube from a 500 μm thick brain slice, which was equal to 18,775 brain cells per sample. With this collection array (**fig. S1**), we divided one brain section (~80 mm^2^) into 310 pixels.

**Fig. 1.**
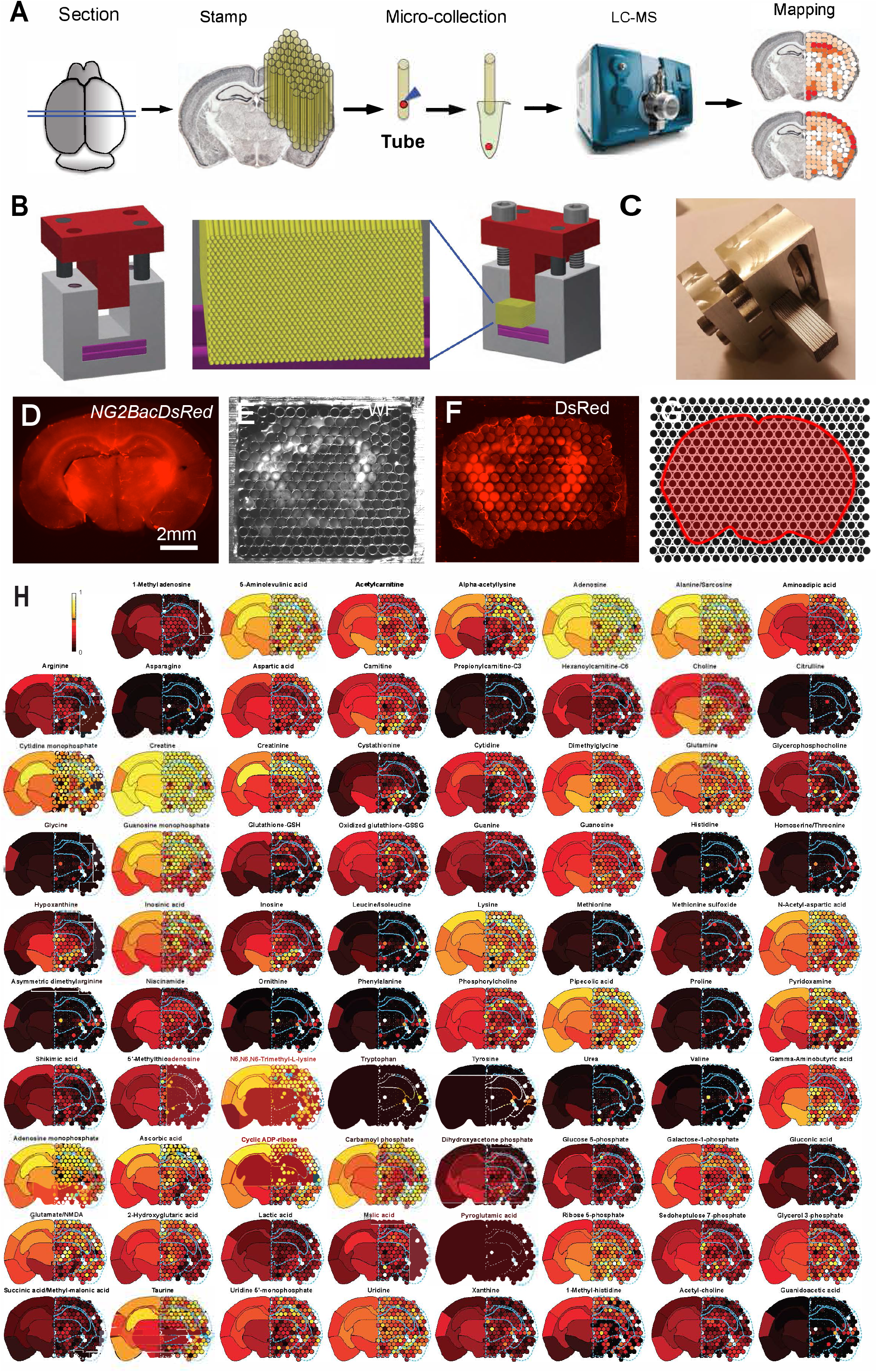
Mapping of 79 metabolites in a mouse brain slice. **(A)** Schematic diagram of spatial metabolomic mapping pipeline. Samples from a brain slice were evenly collected using a microarry collection system (MCA) that we developed. Pixel size and number were determined by the diameter of the microtubes in the MCA. Liquid Chromatography Mass Spectrometry (LC-MS)-based targeted metabolomics were used to analyze the concentration of metabolites from micro-regions (i.e., corresponding to each brain sample collected in a microtube). The values from LC-MS were then applied back to tube locations to make a metabolic atlas of the whole brain slice. (**B)** The design of the MCA for dissecting out fractions of brain tissue. Yellow, steel microtubes; purple, two neodymium-iron-boron magnets; red, the holding bars used to fix all collection microtubes installed with two screws. (**C)** MCA with 450 steel collection tubes on the holding system. (**D**) A brain slice from an adult *NG2DsRedBac* transgenic mouse used for metabolomic analysis. (**E**) White field image of a brain slice (as shown in d) after being collected in the microtubes in the MCA. (**F**) Fluorescent image of the slice that was collected by the MCA. The red fluorescence is DsRed signal from a brain slice of a *NG2DsRedBac* transgenic mouse. The image was illuminated with green fluorescence and then captured under a stereoscope. (**G**) The margin of the brain slice and the locations of all collection tubes. (**H**) Spatial distribution and concentrations of 79 metabolites detected in the mouse brain slice. The whole metabolomic map is combined with the brain macroregion atlas (left hemisphere) and micro-region atlas (right hemisphere). Colors correspond to values in the microregion (i.e., microtube, right hemisphere) or the average value of each microregion located in the corresponding brain macroregion (right hemisphere). The color scale was defined by the maximum and minimum values in all microregions or macroregions. The borders of the different macroregions were determined by corresponding brain sections from Allen Brain Atlas. Color bar represents the relative abundance of a metabolite.

To form a collection array, we designed a holder with a magnet buried in its base, such that it could tightly hold multiple layers of micro-collection tubes (**Fig. 1B**). The magnet was made of neodymium–iron–boron (NdFeB), which is one of the strongest permanent magnetic materials(*20, 21*). Using the holder, we were able to arrange 500–1500 microtubes together and form a collection array for whole brain section collection (**Fig. 1B, C**). We called this whole device the Microtube Collection Array (MCA). With barcodes that we designed for each microtube (**fig. S2**), we could identify the spatial location of each microtube with a unique code; all of these data were then integrated into a virtual image using MATLAB software.

To sample an entire brain section, we stamped a 500-μm adult brain slice mounted on 2% agarose with the MCA. Every tube thus obtained a fraction of agarose with or without a brain sample. Because a block of agarose was also collected in each microtube, it was difficult to tell which microtubes contained a brain sample that entered the tube before the agarose. To overcome this, all brain slices in our measurements were obtained from *NG2DsRedBactg* mice(*22*), in which vascular cells(*20*) and a type of glial cells, NG2 glia, are labeled with red fluorescent protein DsRed (**Fig. 1D-F**). Under green light illumination, the tubes with brain tissues in them were identified through their red fluorescence.

Using our microtube barcode system (**fig. S2**), we could spatially map the original location of each brain sample once we dissembled the tubes after sample collection. The map was drawn from an image of the original shape of the brain section that was taken with a fluorescent stereoscope (**Fig. 1D-F**). With the MCA equipped with OD 508 μm tubes, we divided the whole brain section into about 300 pixels (i.e., samples, **Fig. 1D-G**). After the stamping procedure, all microtubes were removed from the MCA and stored at −80°C. Brain samples in each microtube were moved into a tube with a home-made device (**fig. S1**). After extracting metabolites from the brain sample, we measured 204 metabolites with targeted metabolomic analysis, including 20 amino acids and their derivatives(*23*). These measurements covered most metabolites of the classical cellular metabolic pathways(*24*). Among these 204 metabolites, we reliably obtained good signals from 79 metabolites in each brain sample (**Fig. 1H**).

The relative intensities of individual metabolites for mapping were calculated by normalizing the value obtained from LC/MS for each metabolite individually against the total metabolite value of the entire chromatogram from each sample (i.e. TIC or Total Ion Chromatogram). With the assistance of a customized MATLAB script (see the Supplementary Information) and an Excel VBA macro program based on metabolomic data, the relative intensities were integrated into the corresponding location of the brain map drawn from the brain slice where the sample was collected. The corresponding microregion (i.e., each pixel or sample collected in one microtube of the MCA) of the brain slice was registered back to the corresponding slice map taken with a stereoscope (**Fig. 1D–G, fig. S3a, b**). The value of every pixel of the brain map was assigned using a customized MATLAB program. Two brain slices adjacent to the one used for tissue collection were fixed and then used as a reference for verifying the location of the mapped slice. In addition, we assigned all samples from the brain slice to one of seven major brain regions (we call these the “macroregions”): dorsal cerebral cortex, middle cerebral cortex, ventral cerebral cortex, hippocampus, thalamus, hypothalamus, and white matter/other (**Fig. 1H** and **fig. S3b**). We averaged the relative values of individual metabolites from microregions in each macroregion. The mean values of each macroregion for each metabolite were used for making an average map (**Fig. 1H, fig. S3b**). The color gradient represents the concentration of the metabolites of each macroregion. With this strategy, we made maps for 79 metabolites with high resolution, including 158 microregions of the right hemisphere mirrored by the averaged map of the seven macroregions shown on the left side (**Fig. 1H**).

To identify micro-region clusters with similar metabolomic profiles, we introduced unsupervised non-linear dimension reduction analysis (UMAP)(*13*), a machine-learning algorithm for dimensionality reduction, followed by the Louvain clustering method implemented in the Seurat package(*25*), as used for cell-type clustering in single-cell RNA sequencing analysis(*25, 26*). The results revealed that the samples from one hemisphere grouped into five different clusters (**Fig. 2A**). We mapped the samples of each cluster back to the brain map. Interestingly, most samples from each cluster tended to be located in the same macroregion, which strongly indicated that the microregions from each individual macro-region shared common metabolic pathways (**Fig. 2B**). The observation is also consistent with the results using Principal component analysis (PCA) with samples from the cortex, hypothalamus, thalamus and hippocampus. In PCA analysis, the samples from the same macroregion are clustered together although some metabolites are uniquely enriched in a specific brain macroregion (**fig. S4** and **S5**). For example, we could observe that most samples from cluster 3 belonged to the hippocampus and nearly all samples from cluster 0 were located in the thalamus or hypothalamus. The samples from cortical regions showed much a higher degree of heterogeneity and belonged to three different clusters (clusters 1, 2 and 4). Samples from the dorsal cortical region were significantly different from those in the ventral cortex. For each specific metabolite, we used a UMAP plot to conveniently demonstrate their relative intensities in different clusters. For example, choline and gamma-aminobutyric acid (GABA) were enriched in cluster 0; glycerophosphocholine (GPC) in clusters 0 and 2; and taurine in clusters 1, 2 and 3; while tyrosine and tryptophan were enriched uniquely in cluster 4 (**Fig. 2C**).

**Fig. 2.**
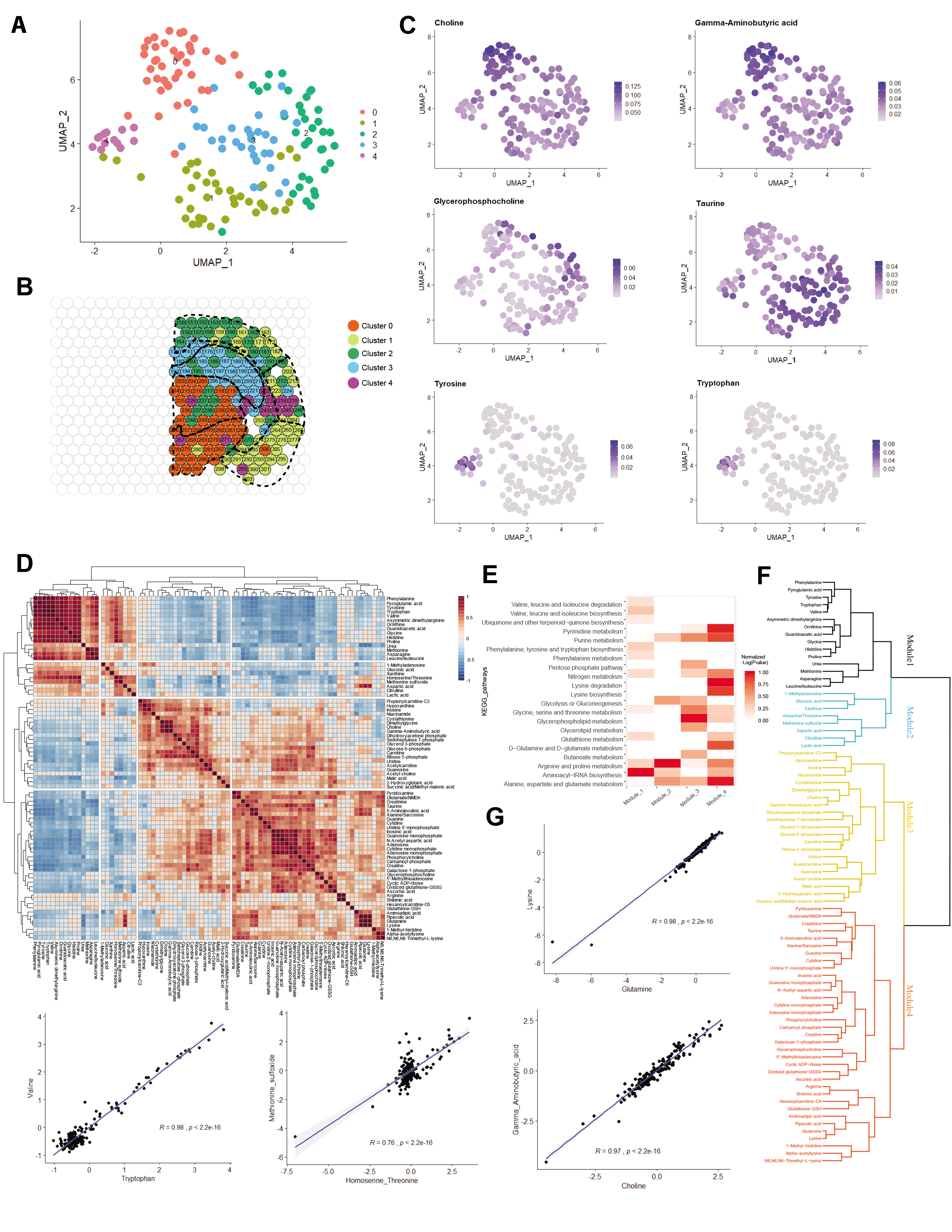
Clustering analysis of metabolomic data from a brain slice. **(A)** UMAP (Uniform Manifold Approximation and Projection) analysis and Louvain clustering of 157 microregions. Five distinct clusters (numbered 0 to 4) were created. Plots produced by Seurat. UMAP visualization of each dot represents a microregion. Clusters of different microregions are color-coded. (**B**) Spatial mapping results of 5 clusters of brain microregions on the microtube arrays. Colors of each well correspond to the dots in (**A**); six representative metabolites enriched from different clusters. (**C**) Metabolite distribution pattern of choline, gamma-aminobutyric acid (GABA, 1^st^ row), glycerophosphocholine, taurine (2^nd^ row), tyrosine and tryptophan (3^rd^ row). In the color bar, purple and gray indicate higher concentration and lower concentrations of a metabolite, respectively. The value of each metabolite was normalized by TIC (total ion value of the entire chromatogram). (**D**) Hierarchical clustering analysis of spatially related metabolites by expression correlation. Data shown as Pearson correlation coefficient. Blue is negative correlation and red is positive correlation. **e**, Pathway analysis results of 4 modules shown in (**D**) (metabolites in each module are marked with dashed lines). Module number listed on axes match the ones listed in the Dendrogram in 2**F**, Hierarchical clustering analysis of metabolite correlation with complete linkage. Detailed demonstration is shown in 2D. (**G**) Scatterplots of linear regression analysis of sub-module metabolites. Four representative pairs of metabolites with close correlation in all microregions from brain section in Fig. 1: glutamine and lysine, tryptophan and valine, homoserine/threonine and methionine sulfoxide, choline and GABA. The letter *r* represents Pearson correlation coefficient. *P* represents the probability value.

There are different metabolic enzymes in different brain cell types, and metabolic enzymes also have distinct expression patterns across different brain regions(*27*). How are different metabolites coupled together to function as a group in cells? A strong spatial correlation among different metabolites indicates that they might be from similar metabolic pathways, or that they are involved in a common cellular signaling pathway or function. To identify spatially related metabolites, we performed Pearson correlation and hierarchical clustering analysis of our spatial metabolic dataset. We observed that 79 metabolites clustered into 4 different modules based on the spatial distribution of their abundance (**Fig. 2D, F**). With these metabolites, we then determined which metabolic pathway(s) were enriched in the brain slice. In Module 1, the enriched metabolic pathway was aminoacyl-tRNA biosynthesis. In Module 2, the enriched pathway was arginine and proline metabolism. In Module 3, glycerophospholipid metabolism was enriched, while a number of pathways including pyrimidine metabolism; purine metabolism; lysine degradation metabolic pathways; alanine, aspartate and glutamate metabolism and lysine biosynthesis metabolism dominated in Module 4 (**Fig. 2E**). From this analysis, we could identify some closely spatially coupled metabolites in the brain, e.g., glutamine and lysine, tryptophan and valine, homoserine/threonine and methionine sulfoxide, choline and GABA (**Fig. 2G**). From the chart, we were able to conveniently determine which metabolites were coupled with others in the adult brain (**Fig. 2F**).

In summary, we have developed an approach and algorithms that allow us to be able to generate a metabolic atlas of a mouse brain section with high resolution (**Fig. 1 & 2**). In addition, we created a micro-region tissue collection system which allowed us to evenly divide a mouse brain section to >300 fractions for targeted metabolomic analysis. With these efforts, we have generated intensity maps for 79 metabolites. Our data have revealed metabolic heterogeneity in major brain regions (**Fig. 2**). This new strategy will make it possible to perform whole-brain metabolic mapping in the near future. The technology can be used for large-scale mapping of brain or tissue metabolomes from large animals.

Mapping the brain metabolome is critical for our understanding of various neurological diseases and physiologies. The same strategy may also be applied to pathological human brain tissue, e.g., resected samples from patients with gliomas or epilepsy, to identify metabolites uniquely enriched in pathological tissues. The subregions of the brain differ with respect to their susceptibility to various pathologies. For example, global ischemia can affect the CA1 region specifically in the hippocampus rather than other brain regions(*28*). Metabolomic mapping may provide researchers with a useful tool to predict the vulnerability of specific brain regions or specific types of neurons to neurological disease. The metabolome atlas provides researchers with a valuable resource to interpret the vulnerability of specific brain regions to various neurodegenerative diseases, and possibly discover specific metabolic biomarkers to define the margin or core of various pathological tissues.

## Author Contributions

W.G., H.Z. and W.S. conceived the project. W.G., W.S. and H.Z. designed the experiments. W.G. established mouse lines, completed slicing, and designed various collection tubes. W.Z. invented the microtube holder for tissue collection. W.G. completed sample collection. W.G., XG., and Fei.C. completed metabolite extraction from all samples. H.Z. performed targeted metabolomic measurement with LC-MS. L.G.Z. and Feng. C. assisted in metabolomic measurement and data interpretation. H.Z., P. H., X.H. J. L. and W.G. completed primary metabolomic analysis from LC-MS. J. L. and W.G. performed all further data analysis in Fig. 1H and Fig. 2 and fig. S5. W. S. and J.L. wrote the code for mapping in Fig. 1H. W.G., H.Z., W.S. and R.J. D. provided reagents. W.G. and J. L. wrote the manuscript. All authors discussed, reviewed and edited the manuscript.

## Acknowledgement

We thank B. Samuels, J. Yu for critical reading of the manuscript. We thank members at Ge and Sun laboratories and other colleagues from CIBR and UTSW CRI for feedback on the work.

---

**fig. S1.**
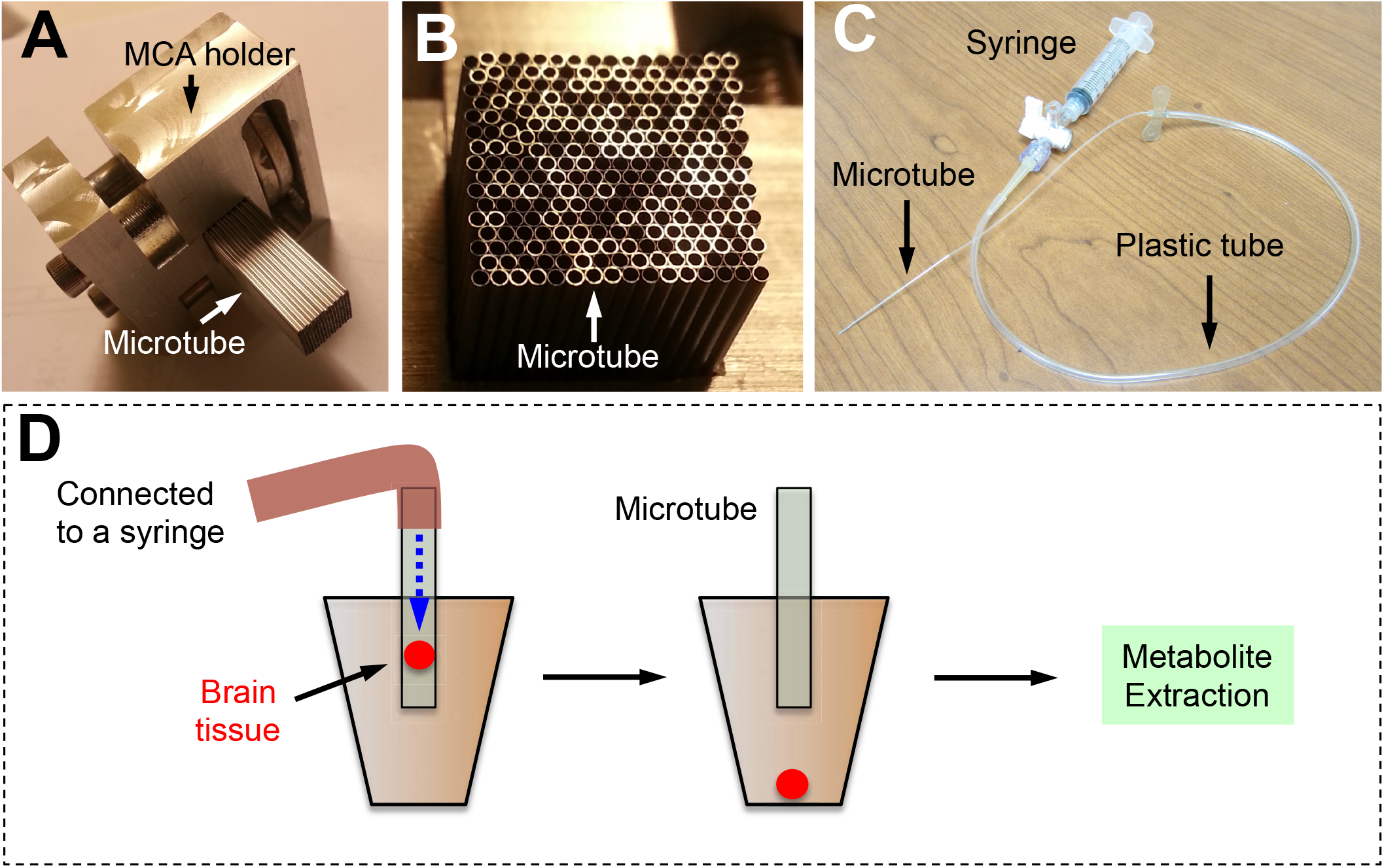
The strategy that we used to collect brain tissue from a microtube into an Eppendorf tube. **(A)** The MCA holder and microtubes. (**B**) An image of the microtube array. (**C**) The microtube was connected to a plastic tube. The plastic tube was linked to a 5ml syringe. (**D**) Positive pressure can be delivered through the syringe and it will move brain tissue from the microtube to the Eppendorf tube. The brain tissue can then be used for metabolite extraction.

**fig. S2.**
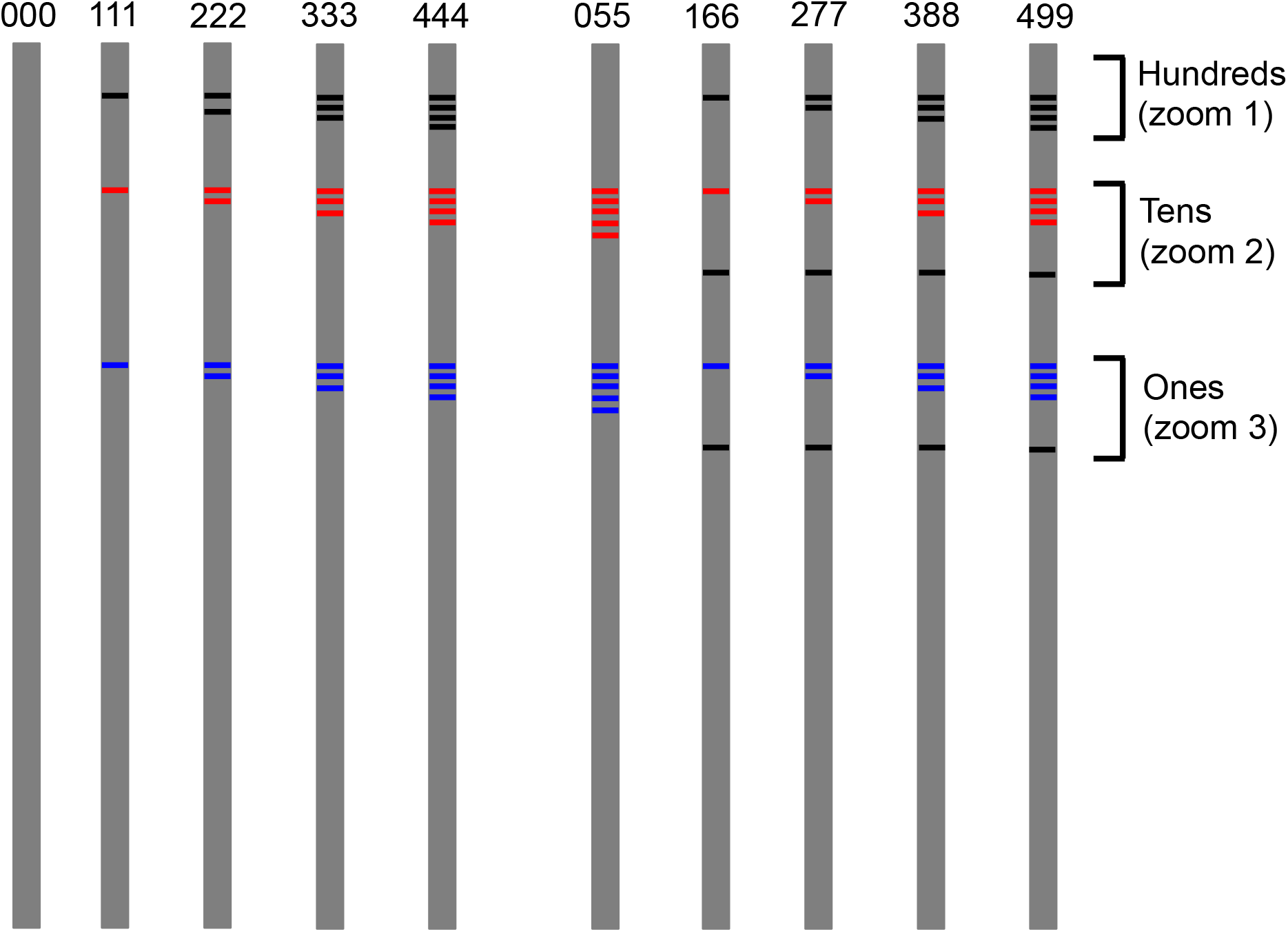
Barcodes that we designed for microtube labeling. The hundreds (100, 200, 300, 400) are marked with black bars at zoom 1. One bar stands for 100. At zoom 2, tens (10 through 90) are marked with red bar(s) together with one black bar. One red bar stands for 10, and one black bar stands for 5. At zoom 3, ones are marked with one blue bar (1–5) together with a black bar. One blue bar stands for 1, and one black bar stands for 5. For example, 8 is marked with three blue bars (i.e., 3) and one black bar (i.e., 5), thus, 3 + 5 = 8.

**fig. S3.**
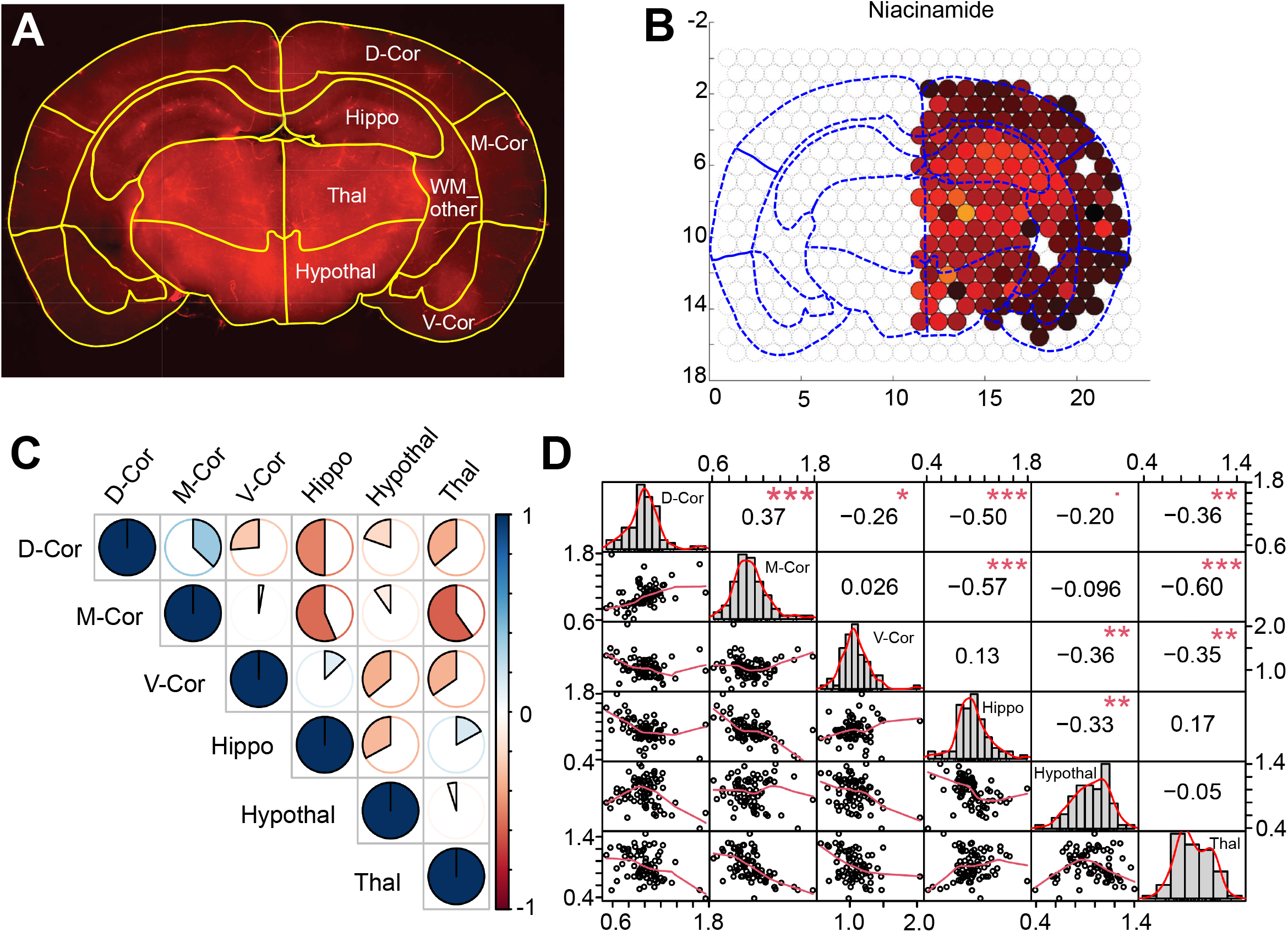
Correlation analysis among different brain macroregions. (**A**) Locations of 7 selected macroregions: D-Cor (dorsal cortex), M-Cor (middle cortex), V-Cor (ventral cortex), Hippo (hippocampus), Thal (thalamus), Hypothal (hypothalamus), and WM_other (White matter and other regions). (**B**) Microtubes in each macroregion in the same brain section. (**C**) Correlation pie plot showing the metabolomic correlations among different macroregions; upper triangular matrix is demonstrated and Pearson method was applied. Positive correlations are displayed in blue and negative correlations in red. Color intensity and circle size are proportional to the correlation coefficients. On the right side of the correlogram, the color bar shows the correlation coefficients and the corresponding colors. (**D**) Correlation chart plot showing the significance of metabolomic correlation among different macroregions. Pearson method was applied. The distribution of each variable is shown on the diagonal. On the bottom of the diagonal, the bivariate scatter plots with a fitted line are displayed. On the top of the diagonal, the value of the correlation plus the significance level are represented by asterisks: *p*-values ≤ 0.001, 0.01, 0.05, 0.1, 1 are represented by “***”, “**”, “*”, “.”, and “ ”, respectively.

**fig. S4.**
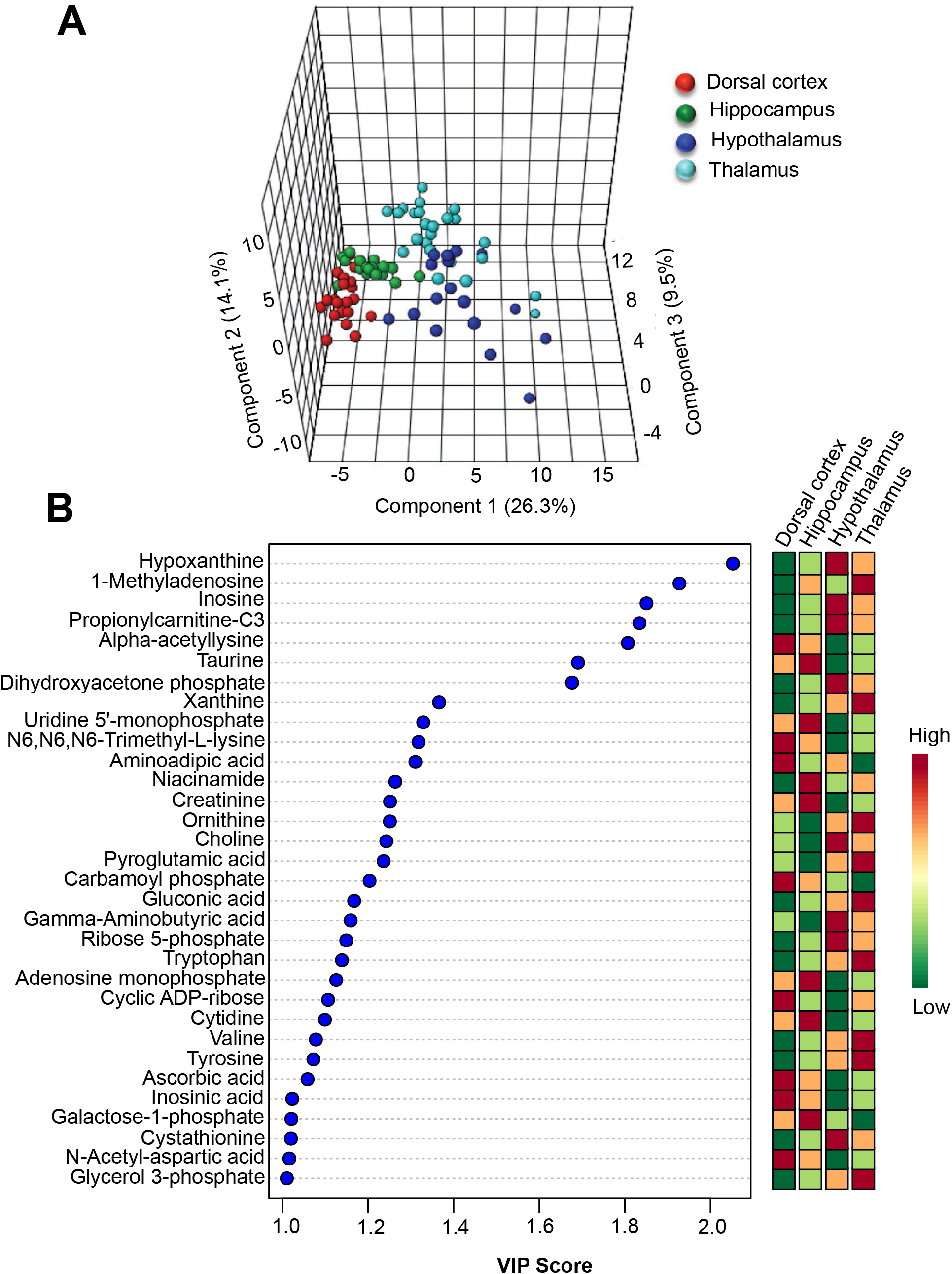
Partial least squares-discriminant analysis (PL-SDA) of metabolites in different brain regions. **(A)** PL-SDA of the 79 metabolites from samples in four different brain macroregions (dorsal cortex, hippocampus, thalamus, and hypothalamus). The top three principal components (PCs) explain 49.9% (26.3%, 14.1% and 9.5%) of the total variance. Each circle indicates an individual sample of one of the four different brain macroregions. As shown in the figure, samples from the different brain macroregions are separated by the PL-SDA. (**B**) VIP analysis of metabolites among samples from the four different brain macroregions. The columns to the right indicate whether the abundance of each metabolite is high (red box) or low (green box) in each brain macroregion. Color bar (bottom left) indicates the scale of VIP values. Warm colors indicate higher concentrations.

**fig. S5.**
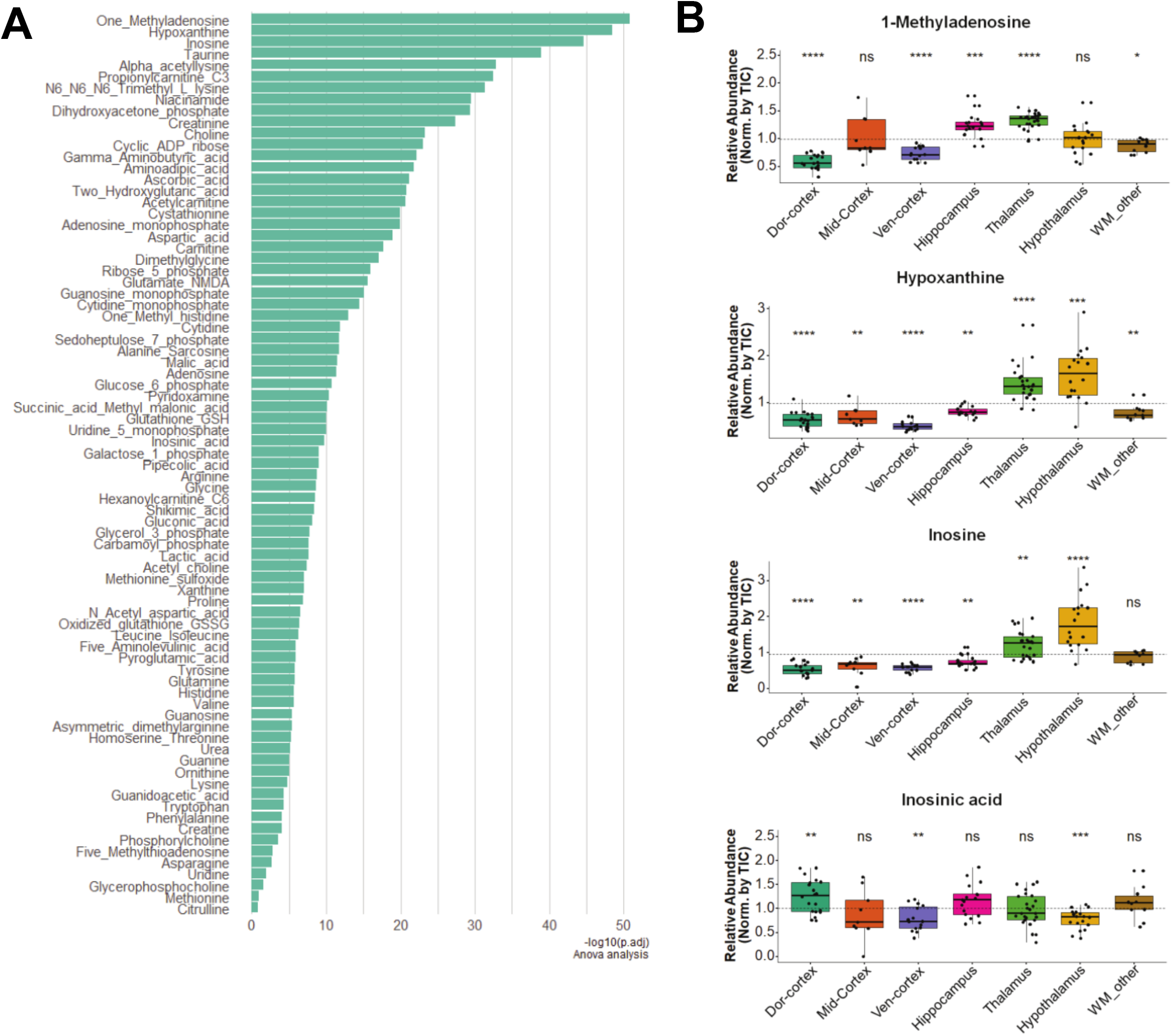
Plotting of all adjusted *p* values obtained through comparing the relative abundance of each metabolite in several different brain macroregions. (**A**) Y axis lists 79 metabolites. The metabolites with lowest adjust *p* values [-log_10_*p* adj] are at the top of the list. (**B-E**) Relative abundance (normalized by TIC, y-axis) of four representative metabolites (1-methyladenosine, hypoxanthine, inosine, and inosinic acid) from 7 brain macroregions. \**p* < 0.05; \*\**p* < 0.01, \*\*\**p* < 0.001, ANOVA. All data are presented as mean ± sem.

## Methods

### Brain slice preparation

Brain slices were prepared as described previously(*29*). The use and care of animals followed the guidelines of the Institutional Animal Care and Use Committee at Chinese Institute for Brain Research and University of Texas Southwestern Medical Center. Adult *NG2DsRedBACtg* mice (P70-100) were anesthetized with isoflurane (5%). After decapitation, the whole brain was dissected rapidly and placed in an ice-cold oxygenated (95% O_2_ and 5% CO_2_) solution containing 119 mM NaCl, 26.2mM NaHCO_3_, 2.5 mM KCl, 1.3 mM MgSO_4_, 2.5 mM CaCl_2_, 1 mM NaH_2_PO_4_, and 11 mM Glucose. Transverse slices (500 μm thick) were cut with a vibratome (model VT-1000S; Leica) and then quickly transferred to a dish with cold agarose for following tissue collection experiments.

### Purification of metabolites from brain tissue

Brain tissue samples were collected in 1.5ml Eppendorf tubes and then 900 μl of ice-cold methanol/80% water (vol/vol) was added. The samples were shaken at 120 rpm on a shaker for 6 h at 4 °C. After centrifugation at 17,000 g for 15 min at 4°C, 850 μl of the supernatant was transferred to a new tube(*23*). Then all samples were evaporated to dryness using a SpeedVac concentrator (Thermo Savant). The samples were then stored in a −80 °C freezer before performing metabolomic profiling analysis.

### Targeted metabolomic measurement of metabolites from brain samples

Metabolites were reconstituted in 50 μl of 0.03% formic acid in analytical-grade water, vortex-mixed, and centrifuged to remove debris. Samples were then injected in randomized order onto a SCIEX QTRAP 5500 liquid chromatograph/triple quadrupole mass spectrometer with electrospray ionization (ESI) source in multiple reaction monitoring (MRM) mode as shown in our previous study(*24*). Separation was achieved on a Phenomenex Synergi Polar-RP HPLC column (150 × 2 mm, 4 μm, 80 Å) using a Nexera Ultra High Performance Liquid Chromatograph (UHPLC) system (Shimadzu Corporation). The mobile phases employed were 0.03% formic acid in water (A) and 0.03% formic acid in acetonitrile (B). The gradient program was as follows: 0-3 min, 0% B; 3-15 min, 0% - 100% B; 15-17 min, 100% B; 17-17.1 min, 100% - 0% B; 17.1-20 min, 0% B. The column was maintained at 35°C and the samples kept in the autosampler at 4°C. The flow rate was 0.5 mL/min, and injection volume 20 μl. Sample analysis was performed in positive/negative switching mode. Declustering potential (DP), collision energy (CE) and Collision Cell Exit

Potential (CXP) were optimized for each metabolite by direct infusion of reference standards using a syringe pump prior to sample analysis. The MRM MS/MS detector conditions were set as follows: curtain gas 30 psi; ion spray voltages 1200 V (positive) and −1500 V (negative); temperature 650°C; ion source gas 1 50 psi; ion source gas 2 50 psi; interface heater on; entrance potential 10 V. Dwell time for each transition was set at 3 msec. Samples were analyzed in a randomized order, and MRM data was acquired using Analyst 1.6.3 software (SCIEX). Chromatogram review and peak area integration were performed using MultiQuant software version 3.0.2 (SCIEX). Each sample was processed identically and randomly, The normalized area values were used as variables for the multivariate and univariate statistical data analysis. The chromatographically co-eluted metabolites with shared MRM transitions were shown in a grouped format, i.e., leucine/isoleucine. All multivariate analyses and modeling on the normalized data were carried out using SIMCA-P (Umetrics, Sweden).

### Targeted metabolomic data analysis

Integrated chromatogram peaks of each metabolite were analyzed with MultiQuant software (SCIEX). The ion intensity was calculated by normalizing single ion values against the total ion value of the entire chromatogram(*30*). The data matrix was input into the SIMCA-P software (Umetrics) by mean-centering and Pareto scaling for subsequent analysis. Both unsupervised and supervised multivariate data analyses (MDAs), including PCA and PLS-DA, were applied as previously reported to classify the samples(*24, 31*). To represent the major latent variables in the data matrix, we generated principal components by MDA and visualized them in a score scatterplot using the online software MetaboAnalyst4.0(*32*). VIP scores > 1 between groups were considered as significantly discriminating.

### Tube atlas and region atlas registration

The outlines of 7 different macroregions of the brain section in Fig. 1 were generated using the “edit shape” function of Microsoft PowerPoint (PPT) and aligned with the real slice image taken under a stereoscope (SMZ18, Nikon). The borders of the different brain macroregions were determined by corresponding positions in the Allen Brain Atlas and two adjacent brain slices.

### Generation of microregion and macroregion atlas

The whole metabolomic atlas was made with two parts, the macroregion atlas for the left hemisphere and the microregion (microtube) atlas for the right hemisphere, using Adobe Illustrator (AI). The outline of the right hemisphere was generated by a customized MATLAB program (see our code in Supplementary Information). Metabolomic data used for microregion heatmap atlas generation was normalized according to the mean value of each metabolite feature (x/mean(x’)) and Min-Max normalization method. The color scale was defined by the maximum and minimum values of all metabolites. The concentration value of each metabolite was normalized to the total ion value of the entire chromatogram. The gradation of the color bar was created by using the MATLAB function “hot” and the colors were filled with a corresponding value from each microregion (i.e., microtube) using the MATLAB function “colormap.” The colors were filled with corresponding values in the tube using the MATLAB function “filledCircle.”

Brain macroregions were outlined by using the “edit shape” function of PPT and aligned with the original slice image from a fluorescent stereoscope. The borders of the different brain macroregions were determined by a corresponding brain section from Allen Brain Atlas. In addition, the region heatmap was evaluated by the average value of the microtubes located in the corresponding brain microregions. A list of the tubes and corresponding brain regions is included in the Supplementary Information. In addition, the color scale was defined by the maximum and minimum values as we did in the microregion heatmap and the colors were filled using a customized EXCEL VBA program with the same color gradation as that used for the microregion heatmap.

### Data Normalization

The TIC method of normalization involves normalizing by the sum of one sample. We applied the log transform function which is the same as in the R package MetaboAnalyst 4.0(*32*) to achieve log transformation and the autoscaling function in the R package MetaboAnalyst 4.0 to achieve z-score standardization.

SumNorm<-function(x) {

1000*x/sum(x, na.rm=T);

}

data <-meta_count[,-80]

data<-t(apply(data, 1, SumNorm)) min.val <-min(abs(data[data!=0]))/10

\# generalize log, tolerant to 0 and negative values

LogNorm<-function(x, min.val){

log2((x + sqrt(x^2 + min.val^2))/2)

^}^

data<-apply(data, 2, LogNorm, min.val);

\# normalize to zero mean and unit variance

AutoNorm<-function(x){

(x - mean(x))/sd(x, na.rm=T);

}

### Boxplot

Statistical significance was determined by analysis of variance (ANOVA). *P* ≤ 0.05 was considered significant. Exact *P* values and the corresponding statistical methods are stated in figure legends where relevant. Statistical analyses were performed using ggplot2 and ggpubr packages implemented in RStudio. All boxplots were generated using ggplot2. Parametric *t*-tests and nonparametric Wilcoxon tests were performed using the stat_compare_means function implemented in ggpubr package.

ANOVA was used to calculate the statistical significance in multiple group comparisons for classified sample numbers in each group. Box and Whisker plots demonstrate the full range of variation (boxes: interquartile range; whiskers: median with minimum – maximum). ANOVA was completed using the “stat_compare_means” function of ggpubr package implemented in Rstudio. Dashed lines represent the average abundance of all samples. We used Tukey’s HSD post hoc test to test the variation pattern of metabolite features across different brain regions. Before using ANOVA, the Shapiro-Wilk normality test was applied and Bartlett’s test was used for homogeneity testing of variance of groups.

### Partial Least Squares – Discriminant Analysis (PLS-DA)

We used Supervised Pattern Recognition PLS-DA Model in Metaboanalyst 4.0 for metabolomic profiling of different brain macro-/micro-regions. As a supervised method, PLS-DA can be used for both classification and feature selection. Cross-validation of the algorithm was used to select an optimal number of components for classification. Two feature-importance measures are commonly used in PLS-DA. The importance of a metabolite is measured by a Variable Importance in Projection score (VIP score). We calculate the VIP score of a metabolite as a weighted sum of the squared correlations between this metabolite and the derived PLS-DA components.

### Hierarchical clustering analysis

Before hierarchical clustering analysis, metabolomic data were processed through normalization by sum, log transformation and auto-scaling (z-score) using the functions (Normsum(); logNorm(); AutoNorm()) in the R package MetaboAnalyst4.0(*32*). The metabolite clustering dendrogram was generated by computing a cross-correlation table for all metabolites using the correlation function cor() implemented in R with Pearson method, and then clustering and plotting using the R hclust() function with complete method. For the heatmap visualization of micro-/macro-region correlation and metabolite-metabolite correlation, Pearson correlations were calculated across all metabolites using linear expression data and grouped via hierarchical clustering using the pheatmap function of the pheatmap R package. Brain macro- and micro-region correlation analysis was performed using Pearson’s R coefficient correlation and the correlation matrix was used to do hierarchical clustering with Euclidean distance and Ward.D’s linkage.

### Statistical analysis with R Package Seurat

Recent progress in single-cell transcriptomics has provided researchers a method by which to cluster cells into cell types based on their expression profiles(*33, 34*). We introduced the advanced dimension reduction techniques and clustering method here to visualize and explain our spatial metabolomic data. We followed a standard procedure as suggested in Seurat’s tutorial (v 3.0.2)(*25*) to analyze our slice microregion metabolomic data, with some parameters and steps modified as stated below. The data processing steps included data normalization/scaling, dimension reduction and unsupervised clustering. Unlike the methods used to process single cell-RNAseq datasets, we did not filter out any features or tubes from our metabolomic data when we created the “SeuratObject” and used the metabolomic dataset as an input to Seurat’s analysis pipeline.

### Data normalization in Seurat pipeline

By default, we used the “LogNormalize” method to normalize the data. This is a global-scaling normalization method that normalizes the metabolites’ abundance measurements for each microregion by the total expression, multiplies this value by a scale factor (we used the default value of 10,000), and log-transforms the result. Before dimension reduction, a standard preprocessing step of linear transformation (‘scaling’, z-score transformation) was applied. These processes are implemented using the NormalizedData and ScaleData functions in Seurat.

### Dimension reduction and unsupervised clustering

Before performing clustering analysis, PCA was performed and the top PCs were used to compute the neighborhood graph. We selected PCs for clustering according to the JackStraw() analysis [similar parameters to those in the Seurat tutorial] and Elbow plot (we used 20 PCs).

We used UMAP to demonstrate the clusters and highlight the relative abundance of each metabolite. Due to the randomness of the UMAP algorithm, we applied setseeds() for reproducible analysis and testing of representation robustness. Louvain algorithm, a modularity-based approach to detect communities implemented in FindClusters() function of Seurat package, was used for unsupervised clustering of microtubes. For the clustering step, we used 20 PCs and the default k parameter of 20 for finding neighbors and used a resolution of 1.2 for finding clusters with the Louvain algorithm, which returned 5 clusters of samples. The cluster IDs of every microtube sample were found after we used the Idents function. Then we traced back every sample to the brain slice with clustering colors to compare the clustering result to the spatial distribution of certain metabolomic clusters.

### Correlation analysis

We used the stat_cor() function implemented in ggpubr R package to analyze and visualize pairwise correlations of selected metabolites. Correlation coefficients with *p*-values are shown in the scatterplot.

### Pathway analysis

The pathway enrichment analysis of differential metabolites between two groups was performed using Metaboanalyst 4.0(*32*)(http://www.metaboanalyst.ca). The *Mus musculus* (mouse; mmu; KEGG organisms abbreviation) pathway library was selected for the analysis and the overrepresentation analysis method used in our study was the hypergeometric test. Significance of enrichment score was evaluated by −Log(p value) result calculated using MetaboAnalyst4.0 Pathway Analysis function module. This result was used to generate the heatmap through min-max normalization using R with the function rescale() and visualized with ggplot2 R package. Only the KEGG pathways enriched in at least one submodule and raw *p* value < 0.05 are listed. Module numbers listed on the x-axis (Fig. 2E) match the ones listed in the dendrogram (Fig. 2F).

### Software information

Our scripts were written in MATLAB, R, and EXCEL VBA.

R Studio v1.1.463; R v3.5.1; MATLAB R2018b

### Code availability

The analysis and visualization code in this study is available from the corresponding author upon reasonable request.

